# Identification of collaborative cross mouse strains susceptible to chlamydial induction of hydrosalpinx

**DOI:** 10.64898/2026.02.06.704472

**Authors:** Yi Wu, Ahmed Mohamed Abdelsalam, Huizhou Fan, Victoria K. Baxter, Xin Sun, Guangming Zhong

**Author notes:** Corresponding authors: Xin Sun, The 3^rd^ Xiangya Hospital of Central South University, 138 Tongzipo Rd, Changsha, Hunan 410013, China., Guangming Zhong, Department of Microbiology, Immunology, and Molecular Genetics University of Texas Health Science Center at San Antonio, 7703 Floyd Curl Drive, San Antonio, Texas 78229, USA.

## Abstract

Sexually transmitted infection with *Chlamydia trachomatis* can cause pathology, such as hydrosalpinx, in the upper genital tract, leading to infertility. To investigate how genetic variation affects chlamydial pathogenicity, we screened five strains of Collaborative Cross (CC) mice for susceptibility to intravaginal infection and hydrosalpinx induction by *C. muridarum*, a mouse-adapted chlamydial species used extensively to reveal chlamydial pathogenic mechanisms. In terms of susceptibility to genital chlamydial infection, the five CC strains fell into two categories: CC011 and CC012 were resistant, while CC037, CC042 & CC080 were susceptible. The resistant strains shed significant levels of live organisms from the genital tract for only ∼2 weeks, whereas the susceptible strains shed for ∼4 weeks. However, the resistant CC012 mice developed the highest level of hydrosalpinx, while the susceptible CC042 mice were fully resistant to hydrosalpinx induction. None of the CC mice developed spontaneous hydrosalpinx in the absence of chlamydial infection. The above results, validated macroscopically and microscopically, indicate no correlation between pathology and genital infection, as observed in other inbred mice. Nevertheless, among the two infection-resistant strains, CC012 developed more severe hydrosalpinx than CC011, and hydrosalpinx was positively correlated with live organism shedding from rectal swabs but not from vaginal swabs, supporting a potential role of gastrointestinal chlamydia in chlamydial genital pathogenicity. The above results lay a foundation for using CC mice to further map the genetic determinants that regulate host susceptibility to chlamydial infection and pathogenicity in the female genital tract.

## Introduction

Sexually transmitted infection with *Chlamydia trachomatis* in the lower genital tract may ascend and cause pathology in the upper genital tract, such as hydrosalpinx, leading to infertility in some women (1, 2). The *C. trachomatis*-induced inflammatory responses are believed to contribute significantly to upper genital pathology (3–5). However, the precise mechanisms remain unknown. The mouse-adapted species *Chlamydia muridarum* has been used extensively to study mechanisms of *C. trachomatis* pathogenesis (6–8), as intravaginal inoculation with *C. muridarum* can cause hydrosalpinx in mice, mimicking the tubal pathology induced by *C. trachomatis* in humans. Hydrosalpinx has been considered a surrogate marker for tubal occlusion and tubal factor infertility (6, 9, 10).

When the *C. muridarum* intravaginal infection mouse model was used to reveal chlamydial pathogenic mechanisms, *C. muridarum* was unexpectedly found to spread to the gastrointestinal tract of the same mouse (11) via hematogenous pathways (12, 13). These findings suggest a complex relationship between chlamydial pathogenicity in the upper genital tract and chlamydial colonization in the genital versus gastrointestinal tracts (14). When a naïve mouse is first exposed to *C. muridarum* in the genital tract, genital chlamydia can both ascend to the upper genital tract and infect oviduct epithelial cells (the 1^st^ hit, required for hydrosalpinx initiation) and then spread to the gastrointestinal tract via hematogenous routes (11–13). The gastrointestinal chlamydia can induce chlamydia-specific profibrotic lymphocytes that are then recruited to the injured oviduct as a 2^nd^ hit to promote hydrosalpinx (15, 16). The promotion of hydrosalpinx by gastrointestinal chlamydia is not due to direct infection of the oviduct cells by the gastrointestinal chlamydia organisms, as the gastrointestinal chlamydia fails to spread to the genital tract (17). However, when a naïve mouse is first fed chlamydial organisms in the gastrointestinal tract, the gastrointestinal chlamydia becomes an oral vaccine that induces protective immunity against subsequent chlamydial infection in the genital tract (18). This finding has led to the development of live-attenuated oral chlamydia vaccines (19–21). Thus, the impact of gastrointestinal chlamydia on chlamydial infection and pathogenicity in the genital tract is context-dependent.

The *C. muridarum* intravaginal infection mouse model has enabled the identification of both chlamydial virulence factors and host immune components that play significant roles in upper genital tract pathology (22–25). Moreover, a comparison of 11 inbred mouse strains has identified a range of susceptibility to hydrosalpinx induction by *C. muridarum* (8). For example, A/J mice are resistant, while CBA/1J mice are susceptible to hydrosalpinx induction by *C. muridarum*. The varied susceptibility to hydrosalpinx induction has been correlated with the kinetics of neutrophil responsiveness to *C. muridarum* infection in the genital tract (26). In response to *C. muridarum* infection, A/J mice recruit neutrophils more rapidly in the genital tract than CBA/1J mice. As a result, the A/J mice prevent the development of the long-lasting hydrosalpinx, although both A/J and CBA/1j mice develop robust pyosalpinx. However, due to significant genetic differences between A/J and CBA/1j mice, identifying the genetic determinants that affect host susceptibility to hydrosalpinx induction by chlamydia has been challenging.

Mice belonging to the Collaborative Cross, or CC, strains were derived from 8 genetically diverse inbred progenitor strains: A/J, C57BL/6J, 129S1/SvImJ, NOD/ShiLtJ, NZO/H1J, CAST/EiJ, PWK/PhJ and WSB/EiJ (27). Each CC strain contains portions of the genomes of the 8 progenitor strains, and each CC strain has been bred to homozygosity, allowing the CC to be used for complex trait analysis. The CC has been extensively used to identify susceptibility to pathogen infections, including viral (28), bacterial (29), and eukaryotic (30) pathogens. The success in using the CC mice to identify genetic determinants that impact host susceptibility to pathogen infection and pathogen-induced pathology relies on the distinct susceptibilities of different CC mouse strains.

To determine whether the genetic variation observed in CC mice translates into differences in susceptibility to chlamydial infection and pathogenicity, we screened five representative CC mouse strains using an intravaginal infection model with *C. muridarum*. We found that the five strains of CC strains fell into two distinct categories in susceptibility to chlamydial infection in the genital tract: The strains CC011 and CC012 were resistant to *C. muridarum* infection with a significant live organism shedding course of only 2 weeks, while CC037, CC042 & CC080 were the susceptible strains, with a 4-week shedding course. However, the infection-resistant CC012 mice developed the highest level of hydrosalpinx, while the infection-susceptible CC042 mice were fully resistant to hydrosalpinx induction. These results suggest that CC mice can be used to map the genetic determinants of host susceptibility to chlamydial infection and pathogenicity. Furthermore, among the two infection-resistant strains, the severity of pathology is positively correlated with levels of live-organism shedding from the gastrointestinal tract but not from the genital tract. Thus, further comparisons among resistant mice may identify novel chlamydial pathogenic mechanisms and correlates of pathogenicity, thereby informing future efficacy evaluations of preventive and interventional strategies.

## Materials and Methods

### 1. Chlamydial organisms and infection of CC mice

The *C. muridarum* organisms (clone G13.32.1 derived from Nigg3) used in the current study were propagated in HeLa cells (human cervical carcinoma epithelial cells, ATCC cat# CCL2.1), purified, aliquoted, and stored as described previously (5, 31). Five strains of female CC mice (10 to 18 weeks old) were purchased from the UNC Systems Genetics Core (https://csbio.unc.edu/CCstatus/index.py?run=AvailableLines.information), including CC011, CC012, CC037, CC042, & CC080. The animal experiments were carried out in accordance with the recommendations in the Guide for the Care and Use of Laboratory Animals of the National Institutes of Health (32). The protocol was approved by the Committee on the Ethics of Laboratory Animal Experiments of the University of Texas Health Science Center at San Antonio.

Each mouse was inoculated intravaginally with 20 μl of SPG (sucrose-phosphate-glutamate buffer) with or without 2 X 10^5^ IFUs of live *C. muridarum* organisms. Five days before intravaginal inoculation, each mouse was injected subcutaneously with 2.5 mg of Depo-Provera (Pharmacia Upjohn, Kalamazoo, MI) to synchronize the estrous cycle and increase susceptibility to chlamydial infection. For the intravaginal inoculation, the inoculum was delivered into the mouse vagina using a 200 μl micropipette tip as described previously (5). The tip was carefully and gently inserted into the mouse’s vagina until slight resistance was felt. Then the pipette plunger was fully pressed to release all of the inoculum, and the tip was removed without releasing the plunger.

### 2. Monitoring live C. muridarum organism recovery from swabs

To monitor live-organism shedding, vaginal swabs were collected on different days after intravaginal infection. Each swab was suspended in 500 µl of ice-cold SPG, vortexed with glass beads, and the released organisms were titrated on HeLa cell monolayers in duplicate as described previously (5). The total number of IFUs per swab/tissue was calculated based on the number of IFUs per field, the number of fields per coverslip, the dilution factors, the inoculation volume, and the total sample volume. An average was calculated from the serially diluted and duplicate samples for each swab/tissue. The calculated total number of IFUs/swab was converted into log_10,_ and the log_10_ IFUs were used to calculate means and standard deviation for each group at each time point.

### 3. Evaluating mouse genital tract tissue pathologies and histological scoring

Mice were sacrificed on day 56 post-infection, and the mouse urogenital tract tissues were isolated. Before removing the tissues from the mouse, we evaluated genital tract gross pathology *in situ* with the naked eye. The severity of oviduct hydrosalpinx was scored based on the following criteria (33): 0, no hydrosalpinx; 1, hydrosalpinx detectable visually but requiring microscopic confirmation; 2, hydrosalpinx clearly visible with naked eyes but the size is smaller than the ovary on the same side; 3, equal to the ovary on the same side; 4, hydrosalpinx larger than the ovary on the same side. The severity of uterine horn dilation was scored based on the following criteria (34): Each horn was divided into 4 equal sections or portions. Based on the number of sections occupied by the dilated areas, the horn was scored from 0 to 4: 0, no dilation; 1, horn with visible dilation, but the area was 1 section; 2, horn with a dilated area of 1 but 2 sections; 3, horn with a dilated area of 2 but 3 sections; 4, horn with a dilated area of 3 sections. The sum of the severity scores from both sides of the same mouse was used as the mouse’s score for subsequent statistical analyses. Mice with any level of identifiable gross pathology were counted as pathology positive. After photographing, the excised tissues were fixed in 10% neutral formalin, embedded in paraffin, and serially sectioned longitudinally (with 5 μm/ section). Efforts were made to include the cervix, both uterine horns and oviducts, as well as luminal structures of each tissue in each section. The sections were stained with hematoxylin and eosin (H&E) as described elsewhere (6). The H&E-stained sections were scored for severity of oviduct dilation based on the following criteria: 0, no significant dilatation; 1, mild dilation of a single cross-section; 2, one to three dilated cross-sections; 3, more than three dilated cross-sections; and 4, confluent-pronounced dilation. Scores assigned to individual mice were calculated into means ± standard errors for each group of animals.

### 4. Immunofluorescence assay

HeLa cells grown on coverslips and infected with chlamydia were fixed and permeabilized for immunostaining as described previously (35, 36). Hoechst dye (blue, Sigma) was used to visualize nuclear DNA. For titrating IFUs from swab samples, a mouse anti-chlamydial LPS antibody (clone# MB5H9, unpublished observation) plus a goat anti-mouse IgG conjugated with Cy3 (red; Jackson ImmunoResearch Laboratories, Inc., West Grove, PA) was used to visualize chlamydial inclusions. All immunofluorescence-labeled samples were observed under an Olympus AX-70 fluorescence microscope equipped with multiple filter sets (Olympus, Melville, NY).

### 5. Statistical analyses

The pathology score data were analyzed with the Wilcoxon rank sum test. The Fisher’s Exact test was used to analyze category data, including the % of mice with oviduct hydrosalpinx. The Spearman’s Correlation was used to analyze relationships between mouse oviduct hydrosalpinx pathology and infection courses in the genital or gastrointestinal tracts. The infection courses were recorded as the IFU recovered from each swab collected on different days after infection. The area under the curve (AUC) was calculated as the sum of all IFUs from a given mouse, each multiplied by the corresponding number of days during which the IFU was recovered. The IFUs at individual time points or as AUCs were compared between groups using the Wilcoxon rank-sum test.

## Results

### 1. The genital tract shedding courses of live organisms reveal distinct susceptibility to chlamydial infection among five different strains of CC mice

Since we have previously shown that genital chlamydia can spread to the gastrointestinal tract to promote chlamydial pathogenicity in the upper genital tract (11–16), we monitored both the genital and gastrointestinal tract shedding courses of live organisms in five strains of CC mice following an intravaginal infection with *C. muridarum* (Fig. 1). Comparison of the genital tract shedding courses revealed that the five groups of CC mice fell into two distinct categories with the CC011 & CC012 strains shedding at significant levels for only 2 weeks, thus defined as resistant strains, while the CC037, CC042, & CC080 strains maintaining significant shedding for at least 4 weeks, thus defined as susceptible strains. When the area-under-curves (AUCs) were compared, the genital shedding course of the resistant CC011 strain was similar to that of the resistant CC012 strain but significantly shorter than those of the three susceptible strains. The difference in the genital shedding courses between the resistant and susceptible strains was mapped to the pronged shedding on days 21, 28, & 35, respectively, from the susceptible strains. Finally, the live organisms shed from the gastrointestinal tract were similar among all mice except the genital-resistant strain CC011. The CC011 mice developed a significantly reduced gastrointestinal shedding course.

**Fig. 1.**
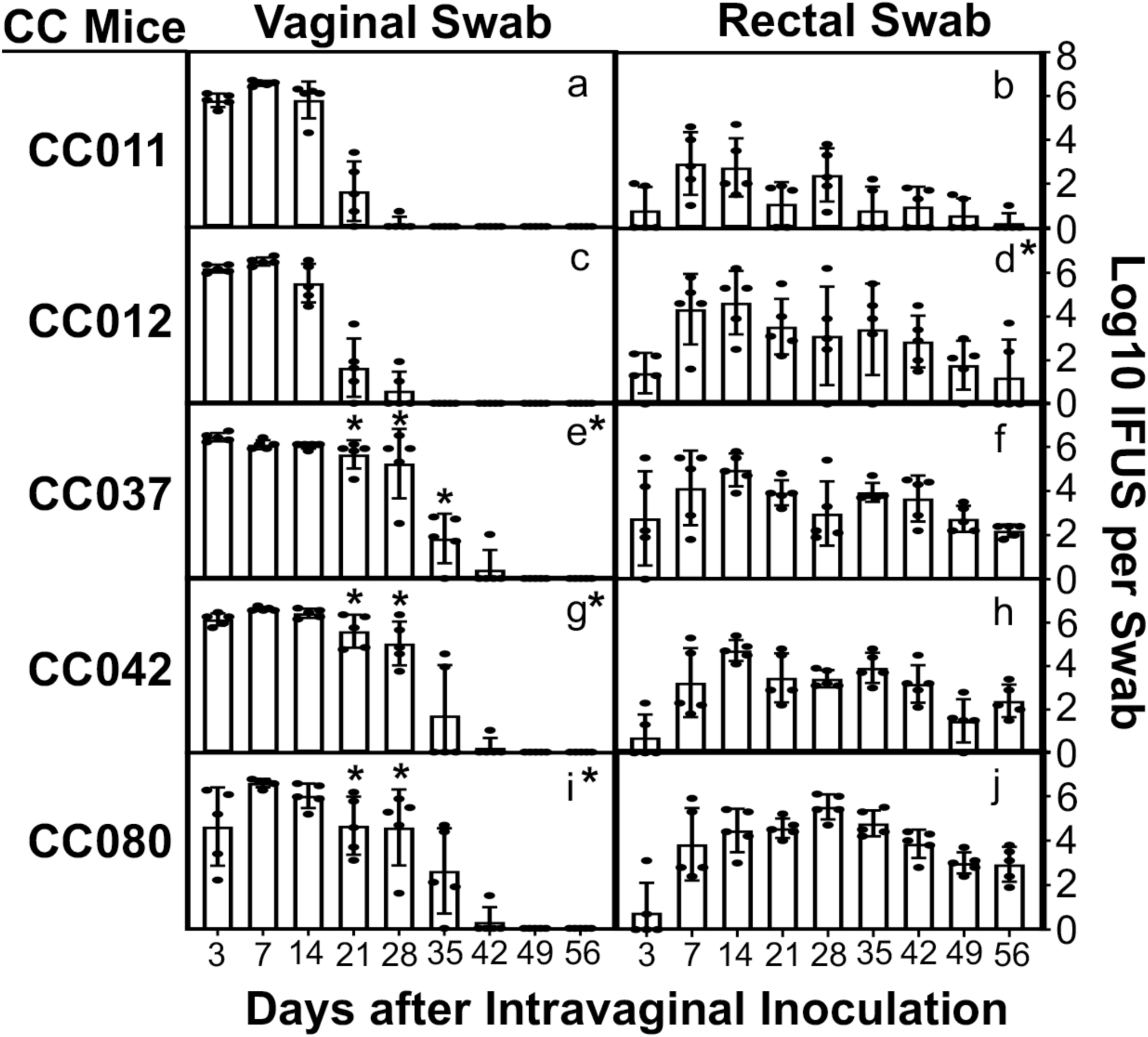
Shedding courses of live chlamydial organisms in five strains of CC mice following intravaginal infection with *C. muridarum*. Five groups of CC mice (as indicated on the left of the corresponding plots, n=5/group) were intramuscularly primed with Depo-Provera for 5 days and then intravaginally inoculated with *Chlamydia muridarum* at a dose of 2 X 10^5^ inclusion-forming units (IFUs) per mouse or SPG buffer alone (data not shown). Vaginal (panels a, c, e, g, & i) and rectal (b, d, f, h, & j) swabs were collected on days 3 and 7, and weekly thereafter, after intravaginal inoculation, as displayed along the X-axis, to monitor live chlamydial organism shedding. The chlamydial burdens were expressed as log_10_ IFUs per swab, as shown along the Y-axis. Note that the five CC mouse strains fall into two distinct patterns in genital shedding courses: short for strains CC011 & CC012, and long for CC037, CC042, & CC080. *p<0.05 (two tails, as the susceptibility of different strains of CC mice to chlamydial infection is unknown), Wilcoxon, area-under-curves (AUCs, a vs. c, e, g, or i, respectively, or b vs. d) or time points (days 21, 28, & 35 from panel a vs. those from panels c, e, g, or i, respectively).

### 2. The mouse susceptibility to hydrosalpinx induction is independent of the genital shedding courses of live chlamydial organisms

To determine whether the genital or gastrointestinal chlamydial shedding courses contribute to chlamydial pathogenicity in the upper genital tract, we have evaluated the gross pathology of the CC mouse genital tracts on day 56 after intravaginal infection. As displayed in Fig.2 and Fig.1S, all CC mice with or without chlamydial infection were sacrificed on day 56 to evaluate genital tract pathology with the naked eye. The severity of uterine dilation and hydrosalpinx was scored according to the criteria described in the Materials and Methods. The scores were compared between CC mice with or without chlamydial infection (Fig. 3). Uterine horn dilation was detected in mice with or without chlamydial infection, while hydrosalpinx was detected only in mice with chlamydial infection. Thus, hydrosalpinx is a more reliable indicator of chlamydial pathogenicity, which is the focus of the subsequent analyses. The genital infection-resistant CC012 strain developed the highest level of hydrosalpinx under the naked eye.

**Fig. 2.**
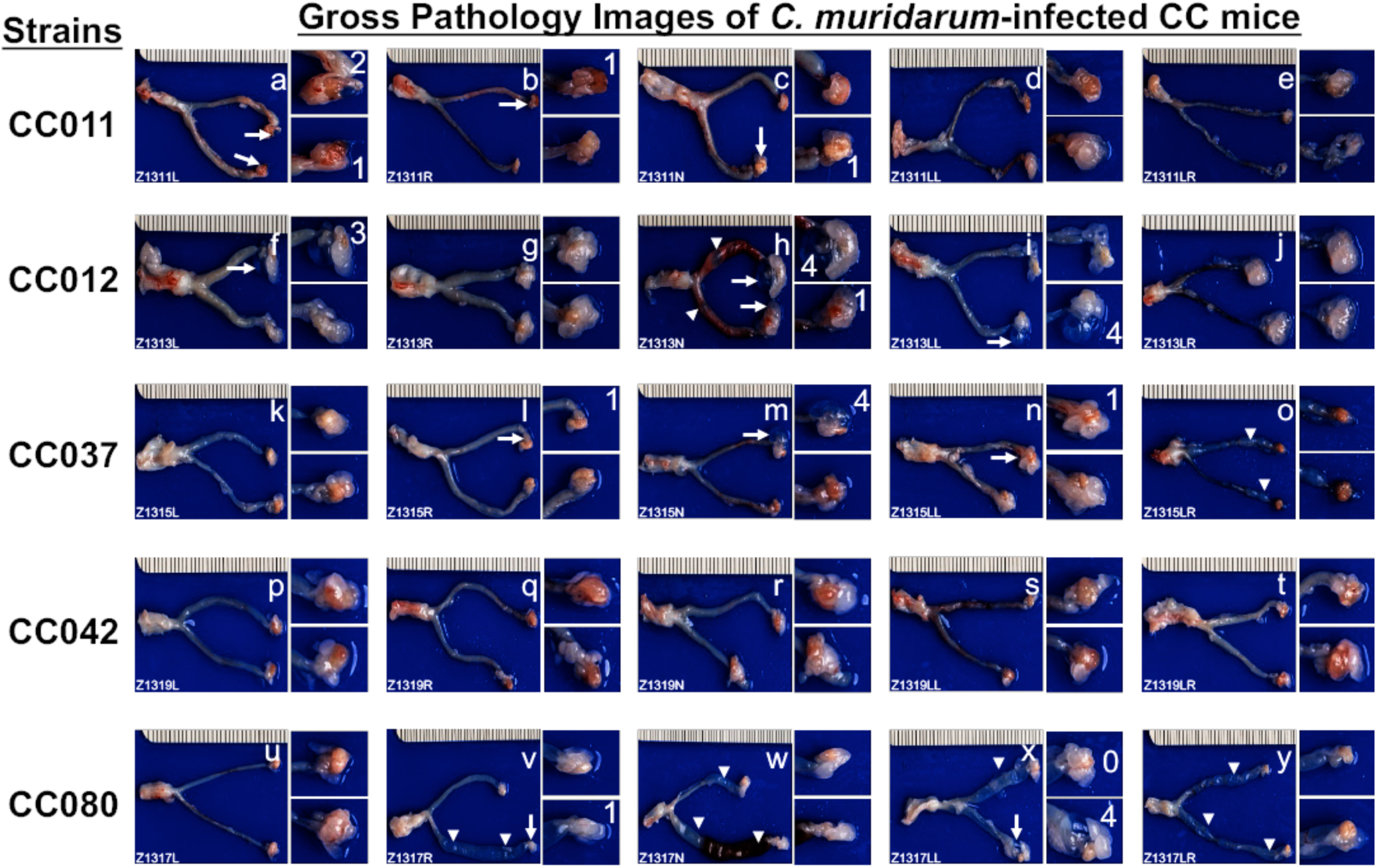
Gross pathology images of the female genital tracts from five strains of CC mice on day 56 after intravaginal infection with *C. muridarum*. The five CC strains of mice, as indicated on the left of the corresponding images (n=5), were intravaginally infected with *C. muridarum* as described in the legend of Fig. 1. On day 56 after infection, all mice were sacrificed to observe genital tract pathology, including uterine horn dilation (white arrowheads) and oviduct hydrosalpinx (white arrows). The image from each infected mouse is presented with the whole genital tract on the left and the magnified oviduct/ovary portion on the right, while the images from buffer-inoculated mice are presented in Supplemental Figure 1. The individual mouse ID is displayed in the lower-left corner of the corresponding image. The severity of each hydrosalpinx was scored according to the criteria described in the Materials and Methods and marked with white numbers in the corresponding images when the score was 1 or higher.

**Fig. 3.**
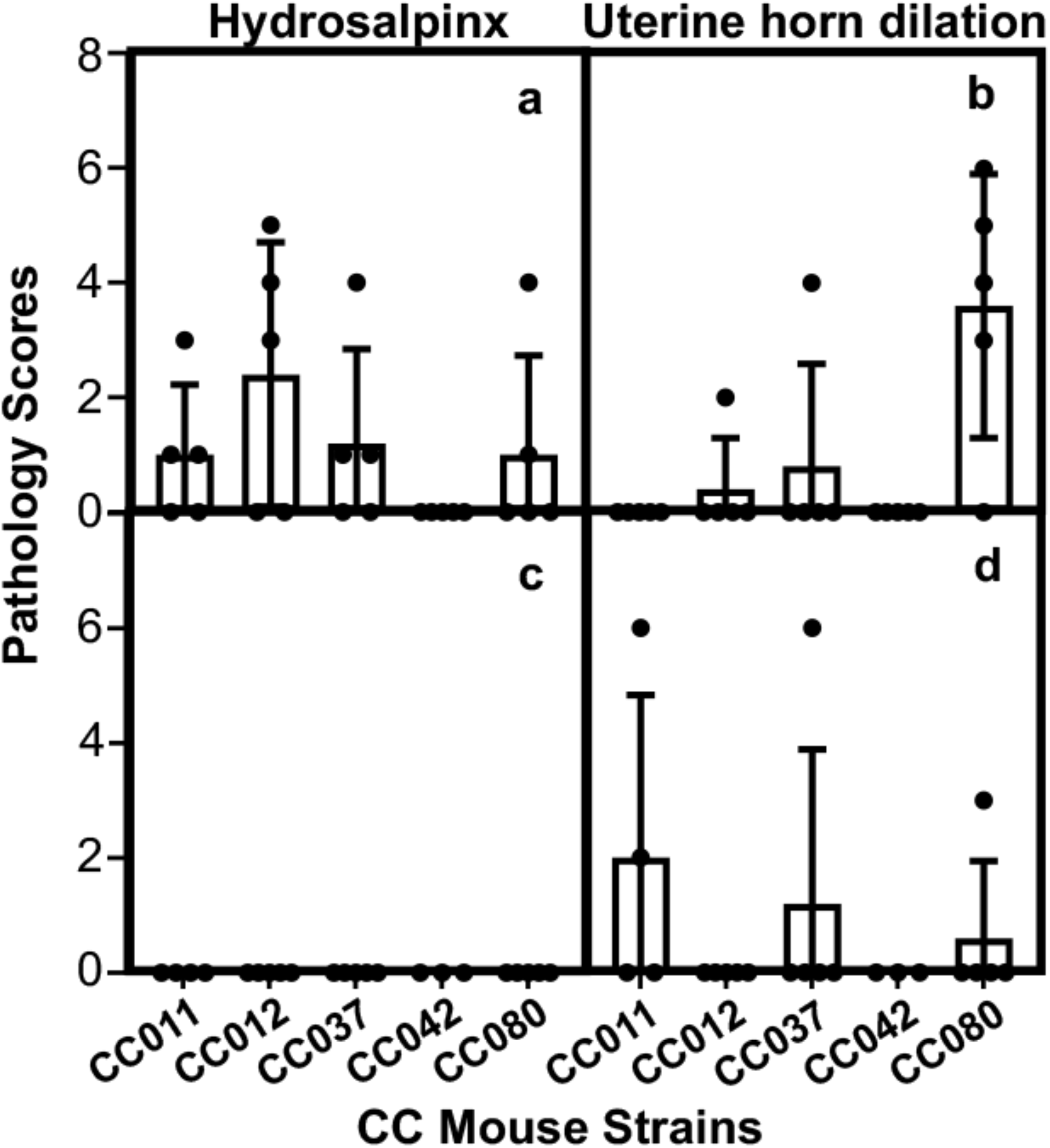
Gross pathology scores of CC mice with or without *C. muridarum* infection. Five strains of CC mice intravaginally inoculated with *C. muridarum* (panels a & b) or SPG buffer alone (c & d), as described in Fig.1 and Fig.1S legends, were sacrificed on day 56 after the inoculation to observe genital tract gross pathology as shown in Fig.2 and Fig.1S. Both hydrosalpinx (a & c) and uterine horn dilation (b & d) were analyzed. The pathology severity scores were compared between CC mice with or without chlamydial infection. Note that hydrosalpinx is only induced by chlamydial infection, while uterine horn dilation occurs spontaneously in CC mice. Wilcoxon, p=0.036 (hydrosalpinx scores between CC012 mice with or without *C. muridarum* infection, assuming chlamydial induction of hydrosalpinx) and p=0.022 (uterine horn dilation scores between CC080 with or without *C. muridarum* infection, assuming chlamydial induction of uterine horn dilation).

### 3. Oviduct dilation observed under microscopy validates the hydrosalpinx observed under the naked eye

Since hydrosalpinx detected in CC mice is dependent on chlamydial infection (see above), and hydrosalpinx is a medically relevant pathology (2, 10, 37), we further validated the hydrosalpinx pathology in CC mice under microscopy (Fig. 4). The same genital tract tissues shown in Fig. 2 and Fig.1S were subjected to sectioning for H&E staining and microscopic observation. The severity of oviduct dilation in CC mice was assessed according to the criteria described in the Materials and Methods. Comparing the oviduct dilation scores between CC mice with or without chlamydial infection revealed that oviduct dilation is dependent on chlamydial infection, and the CC012 mice developed the highest score of oviduct dilation, which was consistent with the highest hydrosalpinx scores in these mice. Consistently, oviduct dilation scores positively correlated with hydrosalpinx scores, with a Spearman’s coefficient of 0.80.

**Fig. 4.**
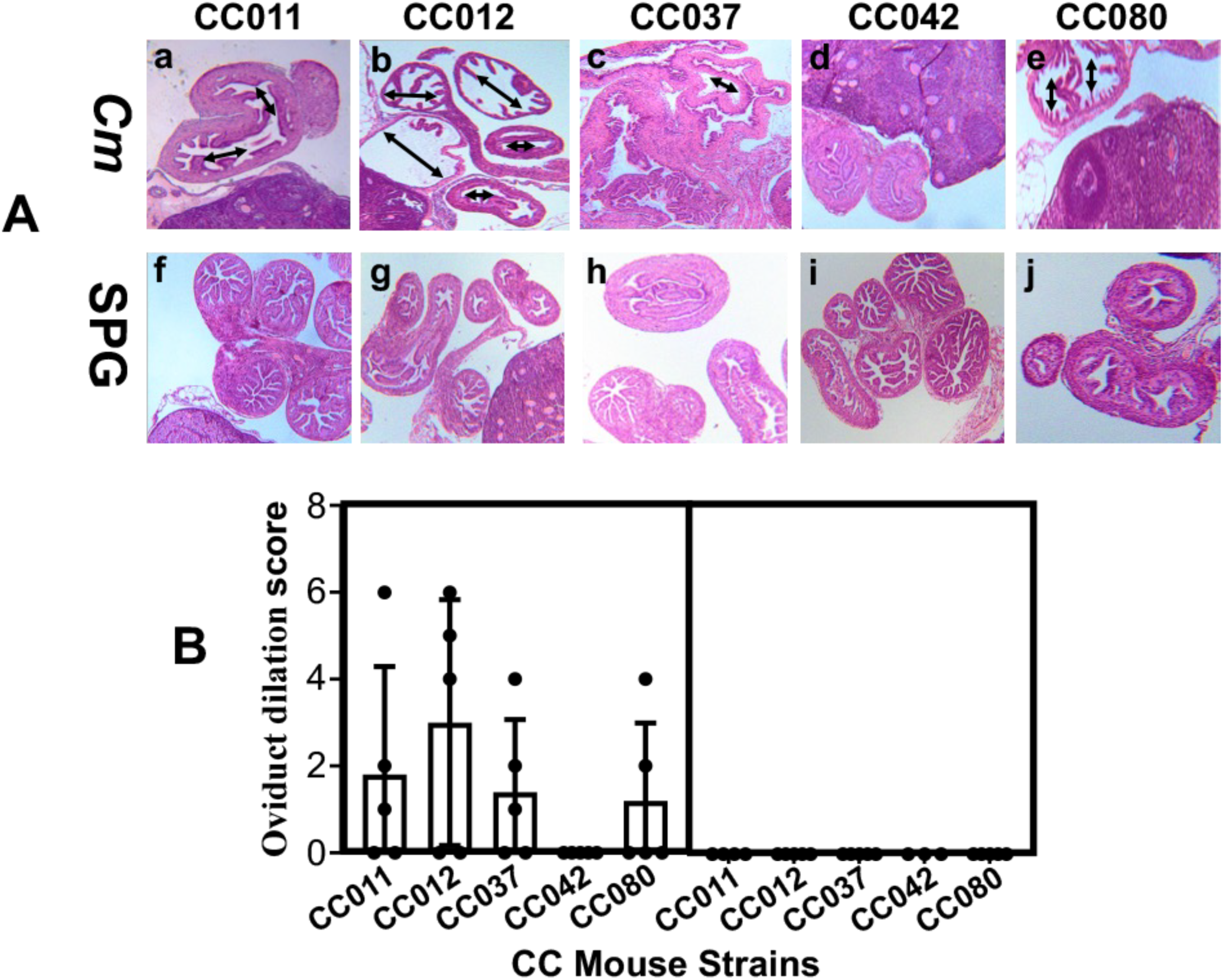
Hydrosalpinx examined under microscopy. The same genital tract tissues shown in Fig. 2 and Fig.1S were subjected to sectioning for H&E staining and microscopic observation. Representative images of the oviduct/ovary portion from each group of mice taken under 4X objective lens are shown. The severity of oviduct lumenal dilation was scored based on the criteria described in the Materials and Methods section. The oviduct lumenal dilation scores were compared between mice with or without chlamydial infection. Note that chlamydial infection induced a significant level of oviduct dilution in CC012 mice. Wilcoxon, p=0.036 (dilation scores between CC012 mice with or without *C. muridarum* infection, assuming chlamydial induction of oviduct dilation).

**Fig. 5.**
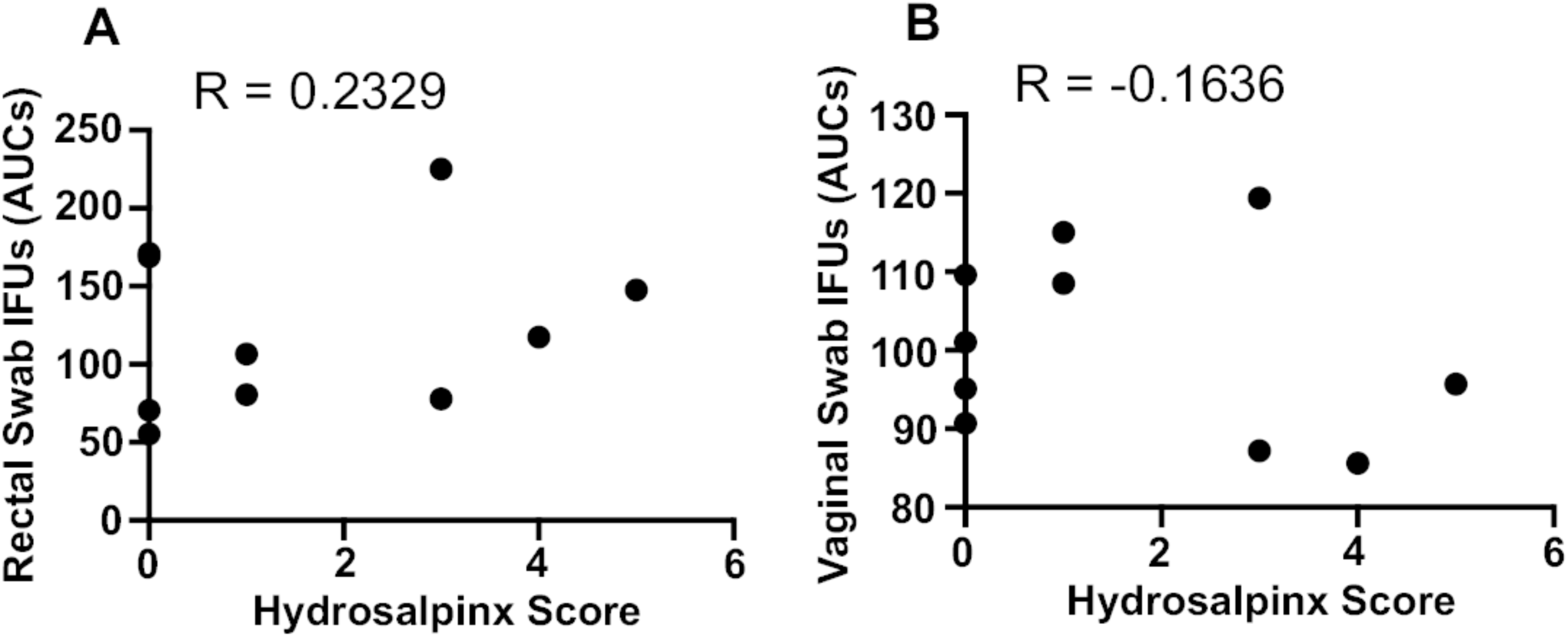
Correlations between hydrosalpinx scores and live organism shedding courses from the genital vs. gastrointestinal tracts of CC mice. The hydrosalpinx severity scores listed in Fig. 3 were correlated with the levels of live chlamydial organism shedding from the gastrointestinal (panel A) versus genital (B) tracts of *C. muridarum*-infected CC011 and CC012 mice (shown in Fig. 1), using Spearman’s correlation (Prism GraphPad, https://www.graphpad.com). Note that the hydrosalpinx displayed a positive correlation with gastrointestinal shedding (R=0.2329) but a negative correlation with the genital shedding (R=-0.1636).

### 4. The upper genital pathology positively correlates with the live chlamydial shedding from the gastrointestinal tract, but not the genital tract

Having evaluated both upper genital tract pathology and live chlamydial shedding in the CC mice, we next determined whether the pathology correlates with the live organism shedding from vaginal or rectal swabs. When all mice were considered, there was no clear correlation. Since the two genital infection-resistant CC mouse strains displayed similar shedding patterns from the genital tract but distinct shedding patterns from the gastrointestinal tract, we analyzed these mice. The severity scores of hydrosalpinx or oviduct dilation positively correlated with live organism shedding from the gastrointestinal tract but negatively correlated with shedding from the genital tract. This finding is consistent with the two-hit model observed in conventional inbred mice, which emphasizes the role of gastrointestinal chlamydial colonization in promoting profibrotic pathology in the upper genital tract (14). Thus, the CC mice can be used to map genetic loci associated with mouse susceptibility to chlamydial induction of hydrosalpinx.

## Discussion

*Chlamydia trachomatis* infection in the lower genital tract may ascend to the upper genital tract, causing pathology such as hydrosalpinx and leading to tubal factor infertility (2, 10, 37). However, the precise mechanism remains unclear. The mouse genital tract infection with *C. muridarum* model has been used to investigate chlamydial pathogenicity (38). CC mice have been used to investigate the pathogenic mechanisms of microbial infections (28–30). In the current study, we used the genital infection model to screen five CC mouse strains for susceptibility to intravaginal infection and hydrosalpinx induction by chlamydia. First, we have found that the 5 CC mouse strains fall into two categories in terms of their susceptibility to chlamydial infection in the genital tract: The infection-resistant strains are CC011 and CC012, with their genital shedding courses of 2 weeks, while the infection-susceptible strains are CC037, CC042 & CC080, with their infection courses for more than 4 weeks. Second, hydrosalpinx is induced in CC mice by chlamydial infection. Still, no CC mice develop hydrosalpinx in the absence of chlamydial infection, indicating that hydrosalpinx is a reliable indicator of chlamydial pathogenicity in CC mice. Third, the genital infection-resistant CC012 mice developed the highest level of hydrosalpinx. In contrast, the susceptible CC042 mice were fully resistant to hydrosalpinx induction, indicating no correlation between the course of lower genital tract infection and upper genital tract pathology. Finally, among the two infection-resistant strains, the hydrosalpinx pathology positively correlated with the spread of vaginal chlamydial organisms into the gastrointestinal tract, indicating that the CC mice mimic the conventional inbred mice in allowing genital chlamydial spreading into the gut (11–13), and gastrointestinal chlamydia may promote chlamydial pathogenicity in the upper genital tract (15, 16). However, the association of gastrointestinal chlamydia with the upper genital pathogenicity is absent from the three susceptible strains, partially due to the lower level of hydrosalpinx among these strains and/or other mechanisms at play. The current study has demonstrated that resistant CC mouse strains may be more suitable for mapping genetic determinants of host susceptibility to chlamydial pathogenicity in the upper genital tract. In comparison, both resistant and susceptible CC mice can be used to map the host genetic determinants in chlamydial infectivity in the lower genital tract.

The identification of the genital chlamydial infection-resistant strains CC011 and CC012, and the susceptible strains CC037, CC042 & CC080, is significant, as this information lays a foundation for further leveraging the CC mouse system to uncover the genetic basis of host susceptibility to chlamydial infection in the genital tract. Intravaginal inoculation was used in the current study, enabling delivery of chlamydial organisms to the ectocervix and the vaginal fornix (a vault-like space formed as the cervix projects into the vaginal canal). This lower genital tract infection model maximally mimics the route of natural infection in women. Thus, using CC mice to identify genetic determinants of susceptibility of the mouse lower genital tract to chlamydial infection may yield clinically relevant information. Both the resistant and susceptible strains maintained similar infection levels during the 1^st^ 2 weeks after intravaginal inoculation, but they differed significantly in genital shedding levels 2 weeks post-inoculation. These observations suggest that the enhanced susceptibility of susceptible strains is due to a delay in mounting an effective adaptive immune response capable of controlling genital infection. This conclusion is consistent with previous observations that adaptive immunity is required to reduce *C. muridarum* shedding two weeks after infection in other inbred mice (22, 39–41). Both chlamydia-specific humoral and cellular immunity may play significant roles (41, 42). Thus, future studies should focus on defining the genetic basis of the differences in both humoral and cellular adaptive immunity between the resistant and susceptible strains of CC mice.

The finding that CC mice develop spontaneous uterine dilation, whereas hydrosalpinx development in CC mice is induced by chlamydial infection, suggests that hydrosalpinx but not uterine horn dilation should be used to evaluate chlamydial pathogenicity in the upper genital tract. This conclusion is consistent with previous findings from other inbred mice. We previously showed that glandular duct dilation causes uterine dilation (34). Although chlamydial infection can promote the development of uterine horn dilation, the promotion is not dependent on the chlamydial plasmid. However, the chlamydial plasmid is required for inducing hydrosalpinx (25, 43). In addition, the *C. muridarum*-induced hydrosalpinx in mice mimics that observed in *C. trachomatis*-infected women under laparoscopy (6, 10). Thus, hydrosalpinx is a more appropriate indicator of chlamydial pathogenicity.

Unexpectedly, *C. muridarum* infection induced the highest level of hydrosalpinx in the genital infection-resistant CC012 mice, while the susceptible CC042 mice were fully resistant to hydrosalpinx induction. This observation indicates a lack of correlation between the course of lower genital tract infection and upper genital tract pathology, which is consistent with our previous findings (8). Intrauterine inoculation with *C. muridarum* induced significant hydrosalpinx in mouse strains that failed to develop hydrosalpinx following intravaginal inoculation, correlating hydrosalpinx with upper, but not lower, genital tract infection. Interestingly, among the two infection-resistant strains, the hydrosalpinx pathology correlatedbetter with chlamydial shedding in rectal swabs than in vaginal swabs, suggesting a role of gastrointestinal chlamydia in chlamydial pathogenicity in the upper genital tract. This assumption is consistent with the two-hit model we proposed based on the observations from conventional inbred strains (14). The two-hit model emphasizes the pathogenic roles of both oviduct epithelial infection by chlamydia resulting from intravaginal inoculation and ascension (the 1^st^ hit), and the profibrotic responses induced by gastrointestinal chlamydia (the 2^nd^ hit). Specifically, genital chlamydial organisms may both ascend along the genital tract to infect oviduct epithelial cells and spread to the gastrointestinal tract. Gastrointestinal chlamydia has been shown to induce CD8+ T cells that promote the induction of hydrosalpinx by the genital chlamydial organisms (16). The stronger correlation of hydrosalpinx with chlamydial recoveries from the gastrointestinal tract than from the lower genital tract among the 2 resistant strains of CC mice suggests that these strains can be used to further define the genetic basis of the 2^nd^ hit. It is estimated that 10 to 15% women with *C. trachomatis* infection in the lower genital tract (if untreated) may develop upper genital tract pathology (http://www.cdc.gov/std/infertility; ref:(2, 10). Similarly, different inbred strains of mice displayed distinct susceptibilities to hydrosalpinx induction by *C. muridarum* infection in the lower genital tract (8). Most *C. trachomatis*-infected women do not develop upper genital tract pathology. Similarly, among the 11 inbred strains we previously studied, only CBA/J and SJL/J mice (representing ∼18% of the 11 strains) were highly susceptible to hydrosalpinx induction by inoculation of the lower genital tract with *C. muridarum*. The high susceptibility of these two inbred strains to hydrosalpinx induction did not positively correlate with vaginal shedding of live organisms in mouse vaginal swabs, suggesting that the host genetic background may contribute to chlamydial pathogenicity in the upper genital tract by directly influencing the quantity and quality of the 2^nd^ hit. As the CC mouse system is designed to reveal the genetic basis of the observed phenotypes, more CC strains should be screened to map host genetic determinants of chlamydial pathogenicity in the upper genital tract.

We are aware of the limitations of the current study. A primary concern is the lack of sufficient CC mouse strains for complete genetic mapping. Nevertheless, the current study is designed to evaluate the feasibility of using CC mice to investigate chlamydial infection biology and pathogenicity, as these mice have not been previously assessed [although we systematically evaluated 11 strains of conventional inbred mice (8)]. The current study not only demonstrated the feasibility but also provided helpful guidance for future CC mouse experimental design. For example, we should expand the resistant strains of CC mice to map host genetic determinants of chlamydial pathogenicity in the upper genital tract. However, both resistant and susceptible CC mice can be used to map the host genetic determinants in chlamydial infectivity in the lower genital tract.

## Supplementary figure

**Fig. 1S.**
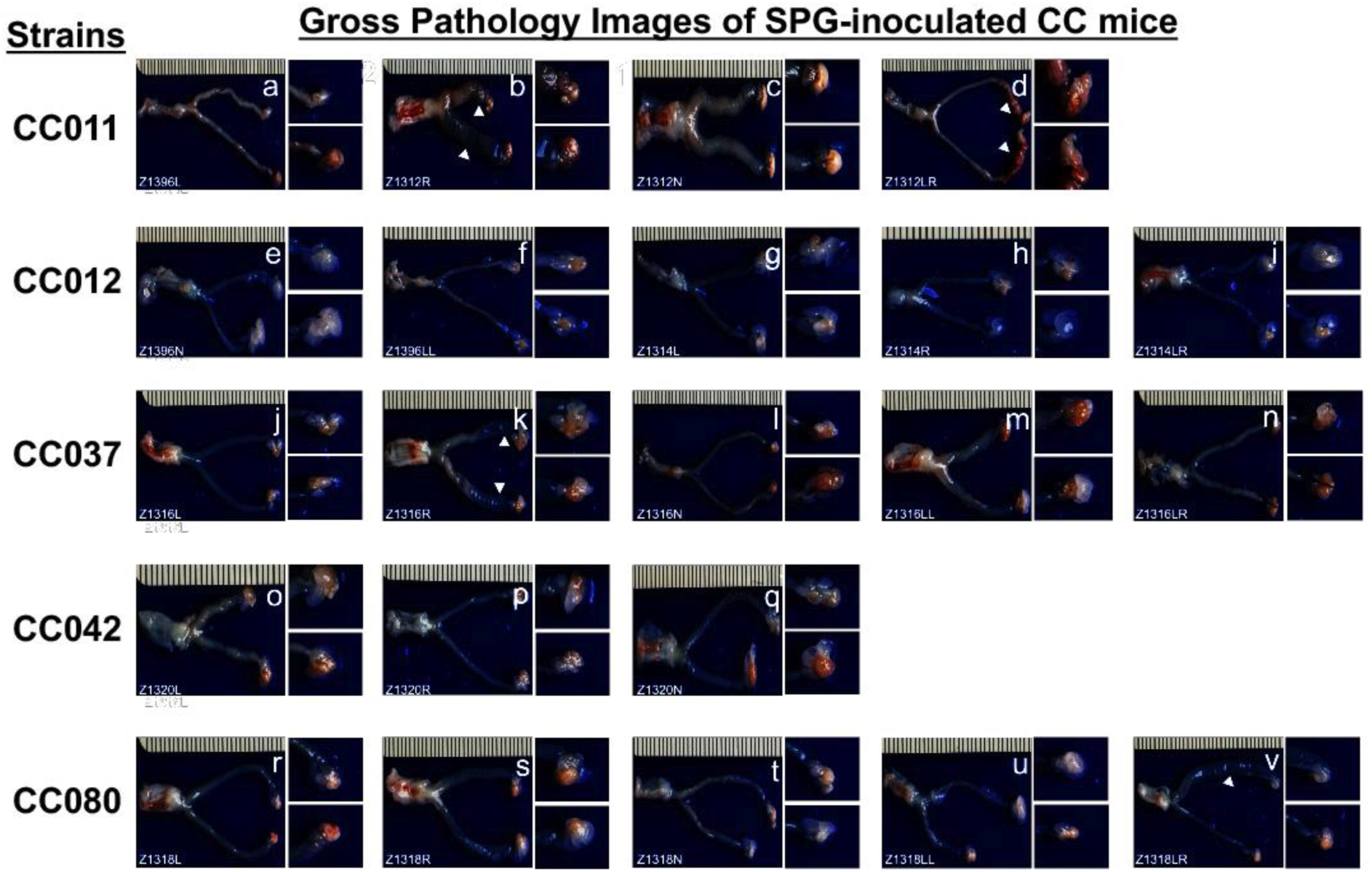
Gross pathology images of the female genital tracts from five strains of CC mice on day 56 after intravaginal inoculation with SPG buffer alone. The five CC strains of mice, as indicated on the left of the corresponding images (n=3 to 5), were intravaginally inoculated with SPG alone as described in the legend of Fig. 1. On day 56 after inoculation, all mice were sacrificed to observe genital tract pathology, including uterine horn dilation (white arrowheads) and oviduct hydrosalpinx (white arrows). The image from each mouse is presented with the whole genital tract on the left and the magnified oviduct/ovary portion on the right. The individual mouse ID is displayed in the lower-left corner of the corresponding image. Note that no significant hydrosalpinx was detected in any of the control CC mice, while uterine horn dilation was detected.

